# Signaling pathways involved in the repression of root nitrate uptake by nitrate in *Arabidopsis thaliana*

**DOI:** 10.1101/2022.10.07.511332

**Authors:** Valentin Chaput, Jianfu Li, David Séré, Pascal Tillard, Cécile Fizames, Tomas Moyano, Kaijing Zuo, Antoine Martin, Rodrigo A. Gutiérrez, Alain Gojon, Laurence Lejay

## Abstract

In *Arabidopsis thaliana*, root high-affinity nitrate (NO_3_^-^) uptake depends mainly on NRT2.1, 2.4 and 2.5, which are repressed by high NO_3_^-^ supply at the transcript level. For *NRT2.1*, this regulation is due to the action of (i) feedback downregulation by N metabolites and (ii) repression by NO_3_^-^ itself mediated by the transceptor NRT1.1(NPF6.3). However, for *NRT2.4* and *NRT2.5* the signaling pathway(s) remain unknown along with the molecular elements involved. Here we show that unlike *NRT2.1, NRT2.4* and *NRT2.5* are not induced in a NO_3_^-^ reductase mutant but are strongly upregulated following replacement of NO_3_^-^ by ammonium (NH_4_^+^) as the N source. Moreover, increasing NO_3_^-^ concentration in a mixed nutrient solution with constant NH_4_^+^ concentration results in a gradual repression of *NRT2.4* and *NRT2.5*, which is suppressed in a *nrt1.1* mutant. This indicates that *NRT2.4* and *NRT2.5* are subjected to repression by NRT1.1-mediated NO_3_^-^ sensing, and not to feedback repression by reduced N metabolites. We further show that key regulators of NRT2s transporters, such as HHO1, HRS1, PP2C, LBD39, BT1 and BT2, are also regulated by NRT1.1-mediated NO_3_^-^ sensing, and that several are involved in NO_3_^-^ repression of *NRT2.1, 2.4* and *2.5*. Finally, we provide evidence that it is the phosphorylated form of NRT1.1 at the T101 residue, which is most active in triggering the NRT1.1-mediated NO_3_^-^ regulation of all these genes. Altogether, these data led to propose a regulatory model for high-affinity NO_3_^-^ uptake in *Arabidopsis*, highlighting several NO_3_^-^ transduction cascades downstream the phosphorylated form of the NRT1.1 transceptor.

**One sentence summary:** Characterisation and identification of molecular elements involved in the signaling pathways repressing NRT2s transporters and root nitrate uptake in response to nitrate.

## Introduction

The nitrogen (N) nutrition of most herbaceous plants relies on the uptake of nitrate (NO_3_^-^), which is ensured in root cells by two classes of transport systems. The High-Affinity Transport System (HATS) is predominant in the low range of NO_3_^-^ concentrations (< 1 mM), whereas the Low-Affinity Transport System (LATS) makes an increasing contribution to total NO_3_^-^ uptake with increasing external NO_3_^-^ concentration (Crawford and Glass, 1998). In all species investigated to date, genes encoding the various transporter proteins involved in either HATS or LATS have mostly been identified in the *NRT2* and *NPF* (formerly *NRT1/PTR*) families, respectively (Nacry et al., 2013; O’Brien et al., 2016). With the exception of agricultural soils, where NO_3_^-^ concentration can rise up to several millimolar after fertilizer application, it is generally assumed that root NO_3_^-^ uptake is mostly determined by the activity of the HATS (Crawford and Glass, 1998; Malagoli et al., 2004). In *Arabidopsis thaliana*, almost all the HATS activity in roots depends on three NRT2 transporters, namely NRT2.1, NRT2.4 and NRT2.5 (Filleur et al., 2001; Kiba et al., 2012; Lezhneva et al., 2014). Under most conditions, NRT2.1 is the main contributor to the HATS (Cerezo et al., 2001; Filleur et al., 2001). However, NRT2.4 and NRT2.5 display a very high-affinity for NO_3_^-^ and are important for taking up this nutrient when present at very low concentration (<50 μM) in the soil solution (Kiba et al., 2012; Lezhneva et al., 2014). Furthermore, unlike NRT2.1 and NRT2.4, NRT2.5 does not require the presence of NO_3_^-^ to be expressed, and is therefore considered crucial for ensuring the initial uptake of NO_3_^-^ as soon as it appears in the external medium (Kotur and Glass, 2015).

Plants have evolved to respond to a challenging environment where NO_3_^-^ concentration in the soil is highly variable in both time and space. Root NO_3_^-^ uptake is strongly regulated in response to changes in the external NO_3_^-^ availability or in the N demand of the whole plant (Crawford and Glass, 1998; Gojon et al., 2009). On the one hand, the HATS activity is quickly stimulated following first NO_3_^-^ supply or re-supply, as a consequence of the so-called primary NO_3_^-^ response (PNR), which is characterized by a rapid induction of *NRT2.1* in the roots shortly (*e.g*., 30 min) after NO_3_^-^ treatment (Tsay et al., 1993; Filleur and Daniel-Vedele, 1999; Lejay et al., 1999; Zhuo et al., 1999; Okamoto et al., 2003). On the other hand, the HATS activity is subjected to a repression exerted on a longer term (*e.g*., several days) by high N status of the whole plant and/or high NO_3_^-^ supply, that down-regulates *NRT2.1, NRT2.4* and *NRT2.5* expression in roots under N satiety conditions (Lejay et al., 1999; Zhuo et al., 1999; Kiba et al., 2012; Lezhneva et al., 2014). This repression is relieved when plants experience N starvation, resulting in a strong increase in HATS capacity that improves NO_3_^-^ uptake efficiency under N limiting conditions (Lejay et al., 1999; Zhuo et al., 1999; Nazoa et al., 2003; Ohkubo et al., 2017; Ota et al., 2020). For *NRT2.1*, repression by high NO_3_^-^ supply is a complex process that requires the concurrent action of two different signaling mechanisms (Krouk et al. 2006). The first one is a feedback downregulation induced by N metabolites that are products of NO_3_^-^ assimilation. This is evidenced by the facts that *NRT2.1* is strongly upregulated in a nitrate reductase (NR) deficient mutant fed with NO_3_^-^ as compared to the wild-type, but is downregulated following ammonium (NH_4_^+^) or amino acids provision (Lejay et al., 1999; Zhuo et al., 1999; Nazoa et al., 2003). The second mechanism is a repression induced by the perception of high external NO_3_^-^ availability by the roots, mediated by the NRT1.1(NPF6.3) transporter acting as a NO_3_^-^ sensor, which is thus referred to as a ‘transceptor’ (transporter/receptor) (Gojon et al. 2011, Maghiaoui et al. 2020). Indeed, repression of *NRT2.1* by high NO_3_^-^ supply is suppressed or strongly attenuated in *nrt1.1* mutants, even in conditions where root N uptake is not reduced by *NRT1.1* deficiency (Munos et al., 2004). This results in the overexpression of *NRT2.1* in normally suppressive conditions (e.g. in NH_4_NO_3_-fed *nrt1.1* plants) along with a lack of stimulation of *NRT2.1* expression by N starvation in *nrt1.1* mutants. Importantly, both repressive mechanisms mediated by NO_3_^-^ and reduced N metabolites signaling need to be active to downregulate *NRT2.1* (Krouk et al. 2006). This explains why high NO_3_^-^ supply fails to lower *NRT2.1* expression in roots of NR-deficient plants (the repression by reduced N metabolites is suppressed), and conversely why high NH_4_^+^ supply also fails to lower *NRT2.1* expression under mixed NH_4_NO_3_ nutrition if the NO_3_^-^ concentration is low, or if NRT1.1 is deficient (the repression by high NO_3_^-^ is suppressed). For *NRT2.4* and *NRT2.5*, the available data do not allow for now to determine whether they obey to the same regulatory model, or not. Both genes are induced by N starvation (Kiba et al., 2012; Lezhneva et al., 2014), but it is not known if this is due to the relief of repression by NO_3_^-^ or reduced N metabolites, or both.

Several genes, mainly transcription factors, have been found to encode regulators of *NRT2.1* repression by high N such as *LBD37-39* (Rubin et al., 2009), members of NIGT1 family (HRS1/NIGT1.4; HHO1/NIGT1.3; HHO2/NIGT1.2; HHO3/NIGT1.1) (Medici et al., 2015; Kiba et al., 2018; Maeda et al., 2018) and members of BTB family namely BT1 and BT2 (Araus et al., 2016). All these regulators are repressors of *NRT2.1* expression in high N conditions, but once again the experiments performed do not allow to distinguish if they are involved in the regulation by reduced N metabolites and/or by high NO_3_^-^. In this context, our study aimed at (i) characterising the regulatory mechanism involved in the repression of *NRT2.4* and *NRT2.5* by high N, and (ii) find regulatory elements involved. By performing experiments on different NH_4_^+^/NO_3_^-^ regimes combined with the analysis of transcriptomic experiments and the use of mutants for the known regulatory elements we were able to clarify the regulation of *NRT2.4* and *NRT2.5* and to refine our knowledge of the NRT1.1 dependent signaling pathway in response to high NO_3_^-^.

## Results

### Regulation of *NRT2.4* and *NRT2.5* by products of NO_3_^-^ assimilation

To discriminate between repression by NO_3_^-^ or by reduced N metabolites for the regulation of *NRT2.4* and *NRT2.5*, we used the *g’4.3* mutant of *A. thaliana*. This mutant is impaired in the first step of NO_3_^-^ assimilation, catalysed by nitrate reductase (NR), and is therefore deficient in reduced N metabolites in the presence of NO_3_^-^ as sole N source. Wild-type and *g’4.3* plants were first grown on a nutrient solution containing NH_4_NO_3_, allowing normal growth of the two genotypes, and then transferred to NO_3_^-^ as sole N source during 24h and 72h (Figure 1). As observed previously, *NRT2.1* level of expression was higher in the roots of the *g’4-3* mutant as compared to the wild-type, especially after transfer to NO_3_^-^ solution (Lejay et al., 1999). In contrast, there is no significant difference between the two genotypes for *NRT2.4* and *NRT2.5* in all conditions (Figure 1). These results indicate that, unlike *NRT2.1*, the regulation of *NRT2.4* and *NRT2.5* does not seem to involve feedback repression by reduced N metabolites produced by NO_3_^-^ assimilation.

**Figure 1.**
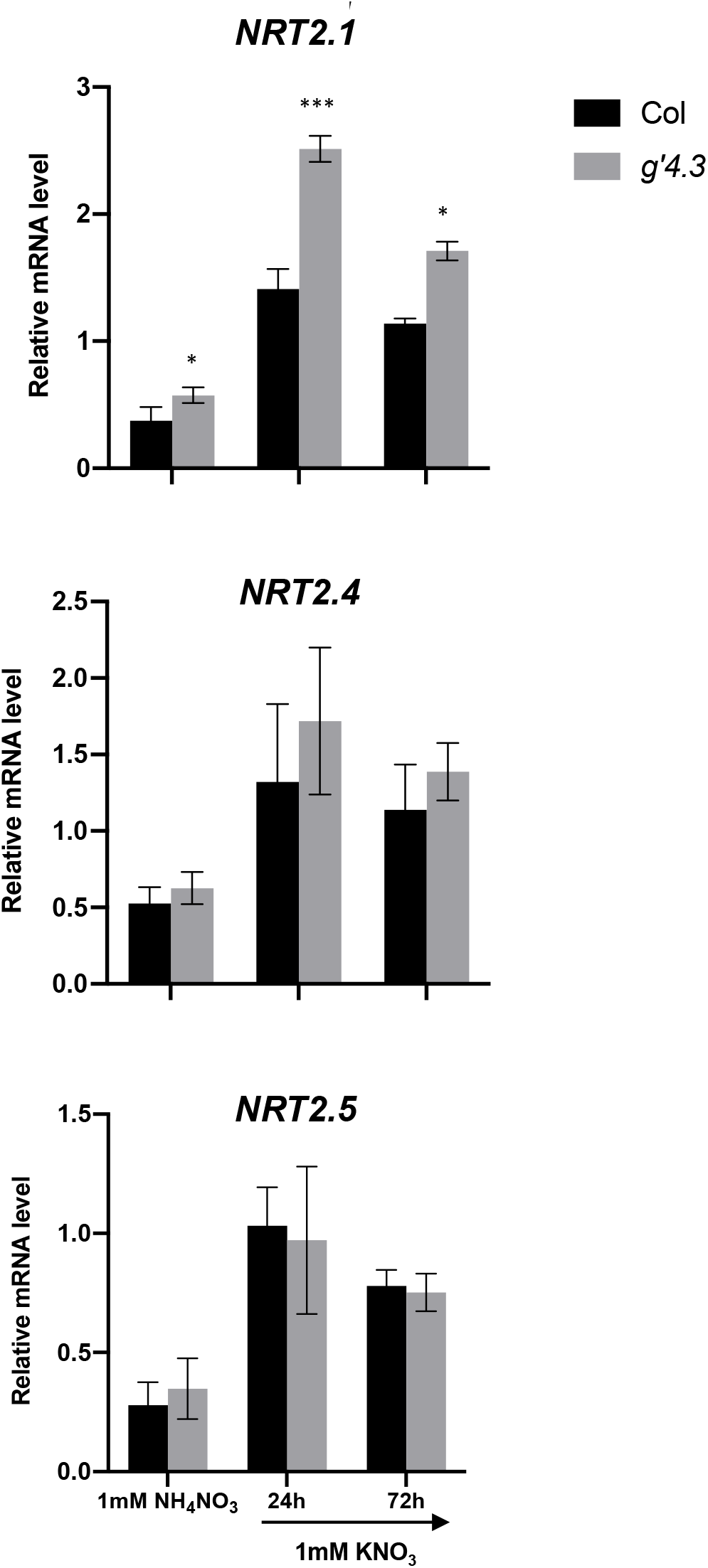
Impact of nitrate reductase mutation (*g’4.3*) on *NRT2.1, 2.4* and *2.5* regulation by the N status. Plants were grown on 1 mM NH_4_NO_3_ for 5 weeks, before being transferred on 1 mM KNO_3_ during 24h and 72h. Roots have been collected to assess *NRT2.1, NRT2.4* and *NRT2.5* mRNA accumulation by RT-QPCR (relative accumulation to *Clathrin* housekeeping gene). Values are means of three biological replicates ± SD. Differences between WT (Col) and the mutant *g’4.3* are significant at \**P* < 0.05, \*\**P* < 0.01, \*\*\**P* < 0.001 (Student’s *t* test).

### Regulation of *NRT2.4* and *NRT2.5* by NO_3_^-^

To investigate the implication of NO_3_^-^ itself in the repression of *NRT2.4* and *NRT2.5*, wildtype plants were grown on 1 mM NO_3_^-^ and transferred during 24h, 48h and 72h on a solution containing 1 mM NH_4_Cl as the sole N source (Figure 2). In those conditions, plants are not starved for N and the only difference is the presence or absence of NO_3_^-^ in the solution. As previously described in other studies, *NRT2.1* is strongly repressed after the transfer of the plants on NH_4_^+^ (Figure 2A) (Munos et al., 2004; Krouk et al., 2006). At the opposite, the expression of both *NRT2.4* and *NRT2.5* transporters is very low on NO_3_^-^ and increased markedly, as soon as 24h, after the transfer of the plants on NH_4_^+^ (Figure 2B). These results confirm that *NRT2.4* and *NRT2.5*, unlike *NRT2.1*, are not repressed by reduced N metabolites and rather suggest that the presence of NO_3_^-^ itself in the nutrient solution is involved in their regulation. In the same experiment, root ^15^NO_3_^-^ influx was measured at two different concentrations. At 50 μM it was correlated with *NRT2.1* level of expression and it decreased when the plants were transferred on NH_4_^+^ (Figure 2A). But when ^15^NO_3_^-^ influx was measured at 10 μM, it tended to increase gradually after 72h on the solution containing NH_4_Cl 1 mM (Figure 2B). It suggests that, when measured at 10 μM, ^15^NO_3_^-^ influx predominantly reflects the activity of *NRT2.4* and *NRT2.5* transporters, especially after 72h on 1 mM NH_4_Cl, when *NRT2.1* expression is the lowest (Figure 2A).

**Figure 2.**
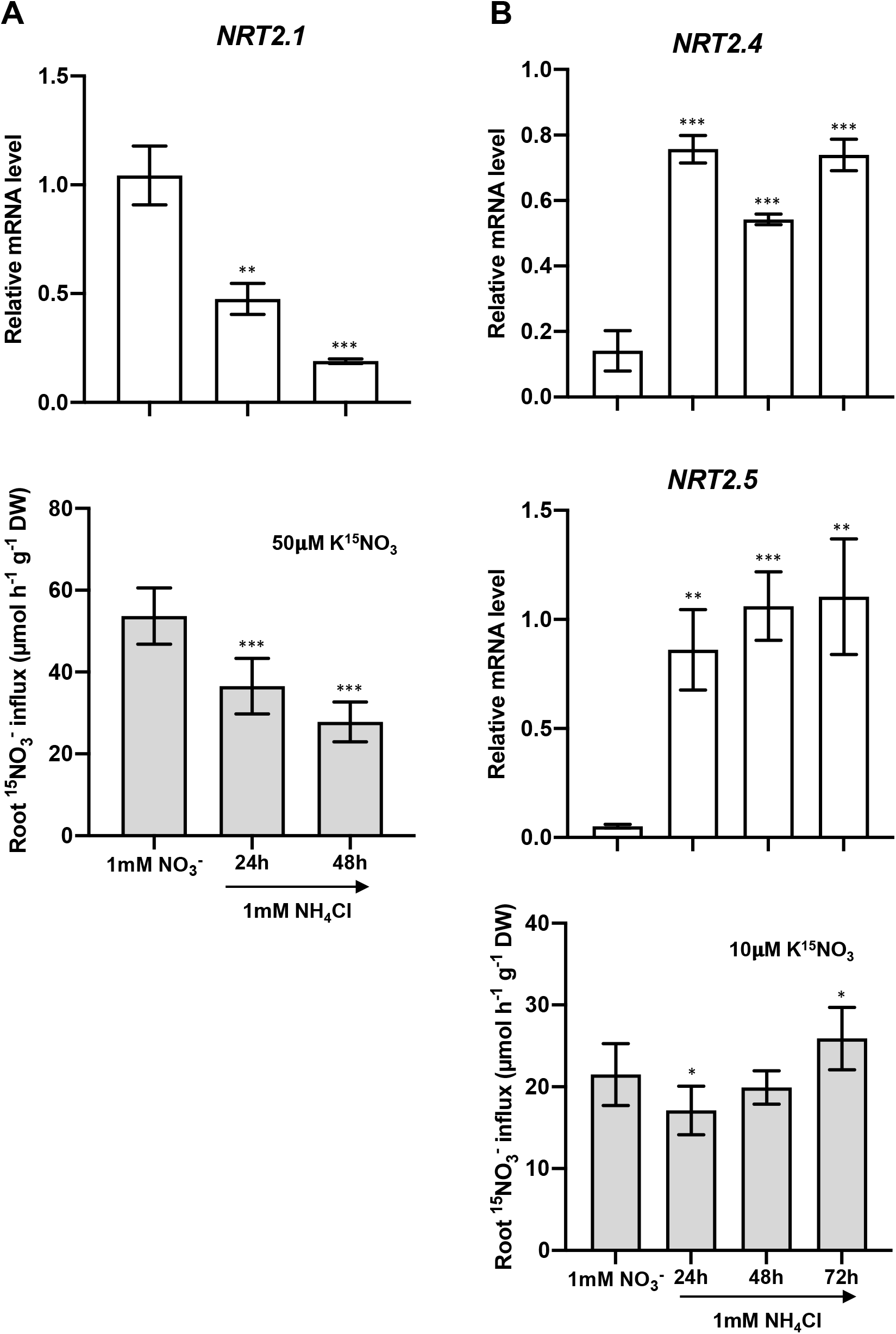
Regulation of *NRT2.1, 2.4, 2.5* expression and of root NO_3_^-^ influx after transfer of the plants from NO_3_^-^ to NH_4_^+^. Plants were grown on 1 mM NO_3_^-^ for 5 weeks before being transferred on 1 mM KNO_3_ during 24h, 48h or 72h. Roots have been collected to assess *NRT2.1, NRT2.4* and *NRT2.5* mRNA accumulation by RT-QPCR (relative accumulation to *Clathrin* housekeeping gene). Values are means of three biological replicates ± SD. Root NO_3_^-^ influx was measured at the external concentration of **(A)** 50 μM ^15^NO_3_^-^ and **(B)** 10 μM ^15^NO_3_^-^’ Plants were treated in the same conditions as for *NRT2s* mRNA level measurements. Values are means of 12 replicates ± SD. Differences between plants on 1 mM NO_3_^-^ and 1 mM NH_4_Cl are significant at \**P* < 0.05, \*\**P* < 0.01, \*\*\**P* < 0.001 (Student’s *t* test).

To confirm these results, suggesting that *NRT2.4* and *NRT2.5* are specifically repressed by NO_3_^-^, were performed experiments adapted from Krouk et al. (2006). Plants were treated with various nutrient solutions that contain 1 mM NH_4_Cl, but differ in KNO_3_ concentration (from 0.1 mM to 10 mM in Krouk et al., 2006). This experimental design revealed that even with an ample supply of NH_4_^+^, which is a strongly repressive condition for *NRT2.1*, this gene was ultimately regulated by NO_3_^-^ signalling, being markedly upregulated when NO_3_^-^ concentration was low (0.1 mM) and repressed when NO_3_^-^ concentration increased up to 10 mM (Krouk et al. 2006). Furthermore, Munos et al. (2004) and Krouk et al. (2006) showed that the repression of *NRT2.1* by high NO_3_^-^ was strictly dependent on NRT1.1/NPF6.3, as the lack of this protein in the mutant *chl1.5* was able to lift it almost totally. To determine if *NRT2.4* and *NRT2.5* are specifically repressed by NO_3_^-^ like *NRT2.1*, we grew wild-type plants and *chl1.5* mutants on 1 mM NH_4_NO_3_ and transferred them during 72h on a mixed solution containing 1 mM NH_4_Cl and 0.1 mM, 1 mM or 5 mM KNO_3_ (Figure 3A). This experiment confirmed that just like *NRT2.1, NRT2.4* and *NRT2.5* are specifically repressed by NO_3_^-^ and that this repression is NRT1.1/NPF6.3-dependent. This is particularly marked for *NRT2.5*, which appeared to be already significantly repressed even at 0.1 mM NO_3_^-^. Interestingly, this regulation has also a strong functional impact since root ^15^NO_3_^-^ influx, measured at 10 μM, was no longer repressed by increasing NO_3_^-^ concentrations in *chl1.5* mutant, compared to wild-type plants (Figure 3A).

**Figure 3.**
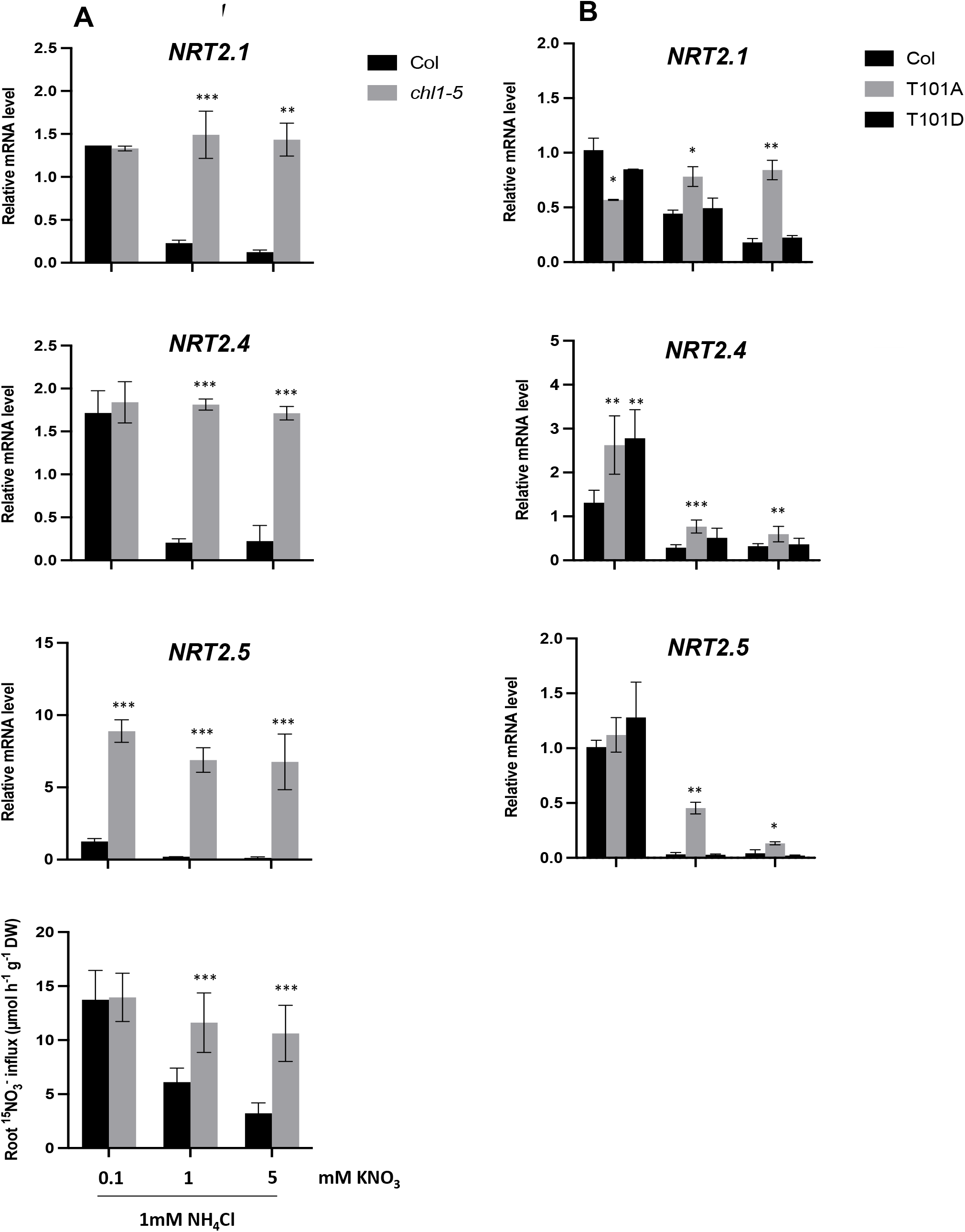
Impact of *NRT1.1* mutations on *NRT2.1, 2.4, 2.5* and root NO_3_^-^ influx repression by NO_3_^-^. Plants were grown on 1 mM NH_4_NO_3_ for 5 weeks before being transferred during 72h on 1 mM NH_4_Cl with 0.1, 1 or 5 mM KNO_3_. Roots of (A) wild type and *chl1-5* knock-out mutant and (B) wild type and T101A and T101D mutants have been collected to assess *NRT2.1*, *NRT2.4* and *NRT2.5* mRNA accumulation by RT-QPCR (relative accumulation to *Clathrin* housekeeping gene). Values are means of three biological replicates ± SD. Root NO_3_^-^ influx was measured at the external concentration of 10 μM ^15^NO_3_^-^ Plants were treated in the same conditions as for *NRT2s* mRNA level measurements. Values are means of 12 replicates ± SD. Differences between WT (Col) and the mutants are significant at \**P* < 0.05, \*\**P* < 0.01, \*\*\**P* < 0.001 (Student’s *t* test).

Furthermore, it was previously shown that phosphorylation of NRT1.1/NPF6.3 at the T101 residue modulates both the transport and signalling activity of the transceptor (Ho et al. 2009). Because repression of *NRT2.1* expression by high NO_3_^-^ was prevented by T101A substitution, which suppresses phosphorylation, while the phosphomimic T101D mutation did not markedly alter it, Bouguyon et al. (2015) suggested that the phosphorylated form of NRT1.1/NPF6.3 is specifically responsible for this repression. We investigated this hypothesis for *NRT2.4* and *NRT2.5*, using NRT1.1/NPF6.3 T101A and T101D mutants. The results showed that T101D mutation was able to phenocopy the repression by high NO_3_^-^ observed in wild-type plants not only for *NRT2.1* but also for *NRT2.4* and *NRT2.5* (Figure 3B). However, compared to *NRT2.1*, T101A mutation was not enough to completely prevent high NO_3_^-^ repression, especially for *NRT2.4*, which was only slightly higher in T101A compared to wild-type plants after 72h on a mixed solution with 1 or 5 mM NO_3_^-^ (Figure 3B). Altogether, these results suggest that like for *NRT2.1*, the phosphorylated form of NRT1.1/NPF6.3 is more specifically active than the non-phosphorylated form for triggering *NRT2.4* and *NRT2.5* repression by NO_3_^-^. However, compared to *NRT2.1*, it seems that repression of *NRT2.5* and mainly that of *NRT2.4* can also be activated by the non-phosphorylated form.

### Molecular elements involved in the repression of *NRT2s* transporters by NO_3_^-^

Following the implication of NRT1.1 phosphorylated form in the repression of *NRT2s* transporter by NO_3_^-^, we used transcriptomic data of Bouguyon et al. (2015) to go further in the characterisation of the signalling pathway involved. In this study, wild-type and several NRT1.1/NPF6.3 mutants were grown *in vitro* on 10 mM NH_4_NO_3_ to find genes regulated by high N concentrations and affected by NRT1.1/NPF6.3 mutation. The analysis of the transcriptomic data revealed, among other things, two particularly interesting clusters (Supplemental Figure 1). A first cluster of 155 genes, whose expression on 10 mM NH_4_NO_3_ is higher in NRT1.1/NPF6.3 mutants compared to wild-type plants and containing *NRT2.1* and *NRT2.4*, as expected from previous results. *NRT2.5* was not part of this cluster likely because the conditions were too repressive to see its expression. A second cluster of 55 genes, whose expression is lower in NRT1.1/NPF6.3 mutants on 10 mM NH_4_NO_3_ compared to wild-type plants. Interestingly, the first cluster contains the protein phosphatase PP2C (At4g32950), called CEPD-induced phosphatase (CEPH), which has been involved in the activation of NRT2.1 by directly dephosphorylating Ser501 of NRT2.1 (Ohkubo et al., 2021). The second cluster contains three members of NIGT1 family (HHO1/NIGT1.3; HHO3/NIGT1.1; HRS1/NIGT1.4), one member of LBD family (LBD39) and one member of BTB family (BT1) (Supplemental Figure 1) (Rubin et al., 2009; Medici et al., 2015; Araus et al., 2016; Kiba et al., 2018; Maeda et al., 2018). These elements have been involved in the repression of *NRT2.1* and *NRT2.4* by N, however it was not known if they play a role in the repression activated by the reduced N metabolites or by NO_3_^-^ itself. To address this question, we tested the impact of *chl1.5* mutation as described above, on plants grown on 1 mM NH_4_NO_3_ and transferred during 72h on a mixed solution containing 1 mM NH_4_Cl and 0.1 mM, 1 mM or 5 mM KNO_3_. We also included *LBD37, LBD38, HHO2/NIGT1.2* and *BT2* in the candidate genes list because they have also been involved in the regulation of *NRT2s* by N (Rubin et al., 2009; Araus et al., 2016; Kiba et al., 2018; Maeda et al., 2018). The results showed that among all these elements, *HHO1, HRS1, LBD37, LBD39, BT1* and *BT2* were induced by NO_3_^-^, while the phosphatase PP2C was strongly repressed by NO_3_^-^ (Figure 4). Altogether, regulations depended on NRT1.1 and more specifically on the phosphorylated NRT1.1 like for *NRT2s* transporters (Figure 4 and Figure 5). This was especially true for both the induction of *HHO1, HRS1, BT1, BT2*, which was significantly impaired on 1 and/or 5 mM NO_3_^-^ in T101A mutant compared to wild-type plants and T101D mutant and for the repression of *PP2C*, which was completely abolished on 1 and 5 mM NO_3_^-^ in T101A mutant (Figure 5). The induction by NO_3_^-^ of both *LBD37* and *LBD39* was less affected by T101A mutation, compared to the other regulatory elements, suggesting that they might be involved in another signalling mechanism dependent on NRT1.1 (Figure 5).

**Figure 4.**
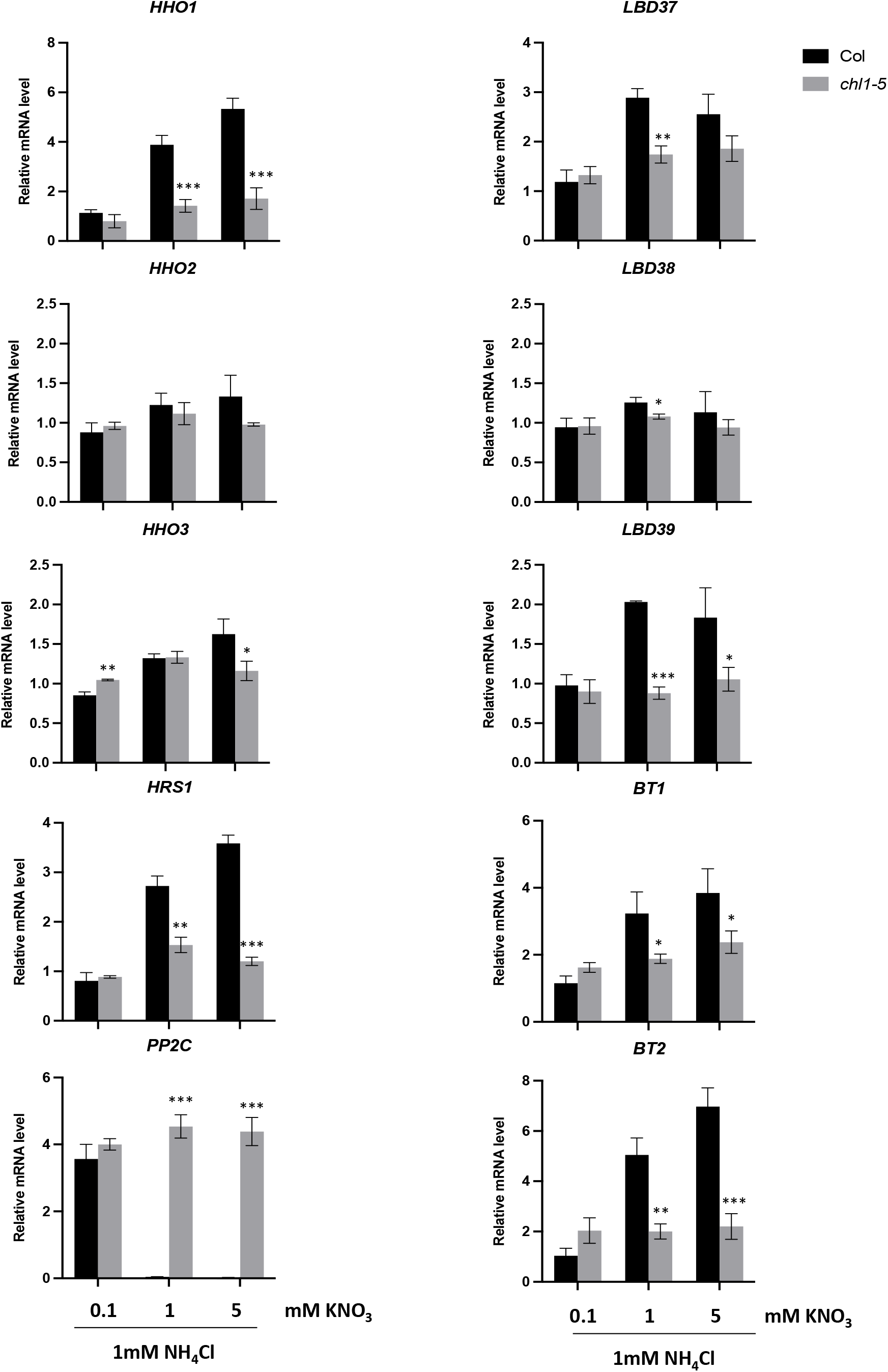
Impact of *nrt1.1* mutation (*chl1.5*) on *HHO1, HHO2, HHO3, HRS1, LBD37, LBD38, LBD39, BT1* and *BT2* regulation by NO_3_^-^. Plants were grown on 1 mM NH_4_NO_3_ for 5 weeks before being transferred during 72h on 1 mM NH_4_Cl with 0.1, 1 or 5 mM KNO_3_. Roots have been collected to assess mRNA accumulation by RT-QPCR (relative accumulation to *Clathrin* housekeeping gene). Values are means of three biological replicates ± SD. Differences between WT (Col) and the mutant *chl1.5* are significant at \**P* < 0.05, \*\**P* < 0.01, \*\*\**P* < 0.001 (Student’s *t* test).

**Figure 5.**
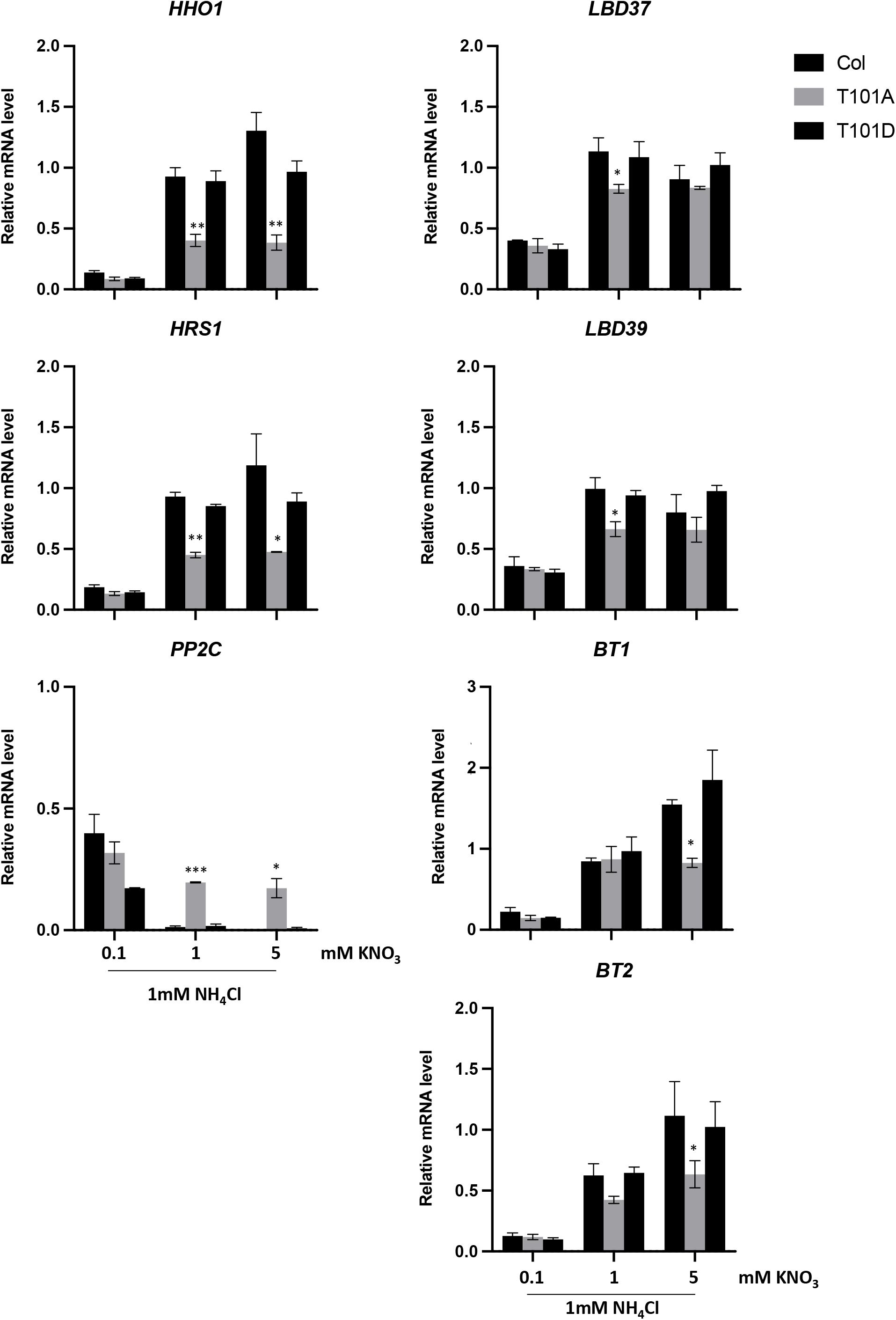
Impact of *NRT1.1* mutations (T101A, T101D) on *HHO1, HRS1, LBD37, LBD39, BT1, BT2* and *PP2C* regulation by NO_3_^-^. Plants were grown on 1 mM NH_4_NO_3_ for 5 weeks, before being transferred during 72h on 1 mM NH_4_Cl with 0.1, 1 or 5 mM KNO_3_. Roots have been collected to assess mRNA accumulation by RT-QPCR (relative accumulation to *Clathrin* housekeeping gene). Values are means of three biological replicates ± SD. Differences between WT (Col) and the mutant *chl1.5* are significant at \**P* < 0.05, \*\**P* < 0.01, \*\*\**P* < 0.001 (Student’s *t* test).

### Impact of the regulatory elements on the regulation of root NO_3_ uptake

To determine the impact of the regulatory elements identified above on the repression of root NO_3_^-^ uptake activity by NO_3_^-^, we measured ^15^NO_3_^-^ influx at 10 μM using the double mutants *hh* (for *HHO1* and *HRS1*), *lbd37/lbd39* and *bt1/bt2*, in the same experimental set up as above, with plants grown on 1 mM NH_4_NO_3_ and transferred during 72h on a mixed solution containing 1 mM NH_4_Cl and 0.1 mM, 1 mM or 5 mM KNO_3_. The results showed that the repression of root NO_3_^-^ influx was only affected in the double mutants *hh* and *bt1/bt2* (Figure 6). Compared to wild-type plants, NO_3_^-^ influx was significantly higher in both double mutants after transfer on increasing NO_3_^-^ concentration at 1 and 5 mM (Figure 6). Conversely, no differences were observed between the double mutant *lbd37/lbd39* and wild-type plants. However, it should be noted that the lack of *HHO1/HRS1* or *BT1/BT2* did not completely prevent NO_3_^-^ repression of root NO_3_^-^ uptake (Figure 6). As expected, the mis-regulation of root NO_3_^-^ uptake activity in the double mutant *hh* was correlated with a complete lack of repression by NO_3_^-^ of *NRT2.1* and a significant higher expression of *NRT2.4* and *NRT2.5* on 1 and 5 mM NO_3_^-^ compared to wild-type plants (Figure 7). But surprisingly, repression of *NRT2.1, NRT2.4* and *NRT2.5* was the same in the double mutant *bt1/bt2* compared to wildtype plants and could not explain the higher level of root NO_3_^-^ uptake activity observed in the double mutant (Figure 6 and Figure 7).

**Figure 6.**
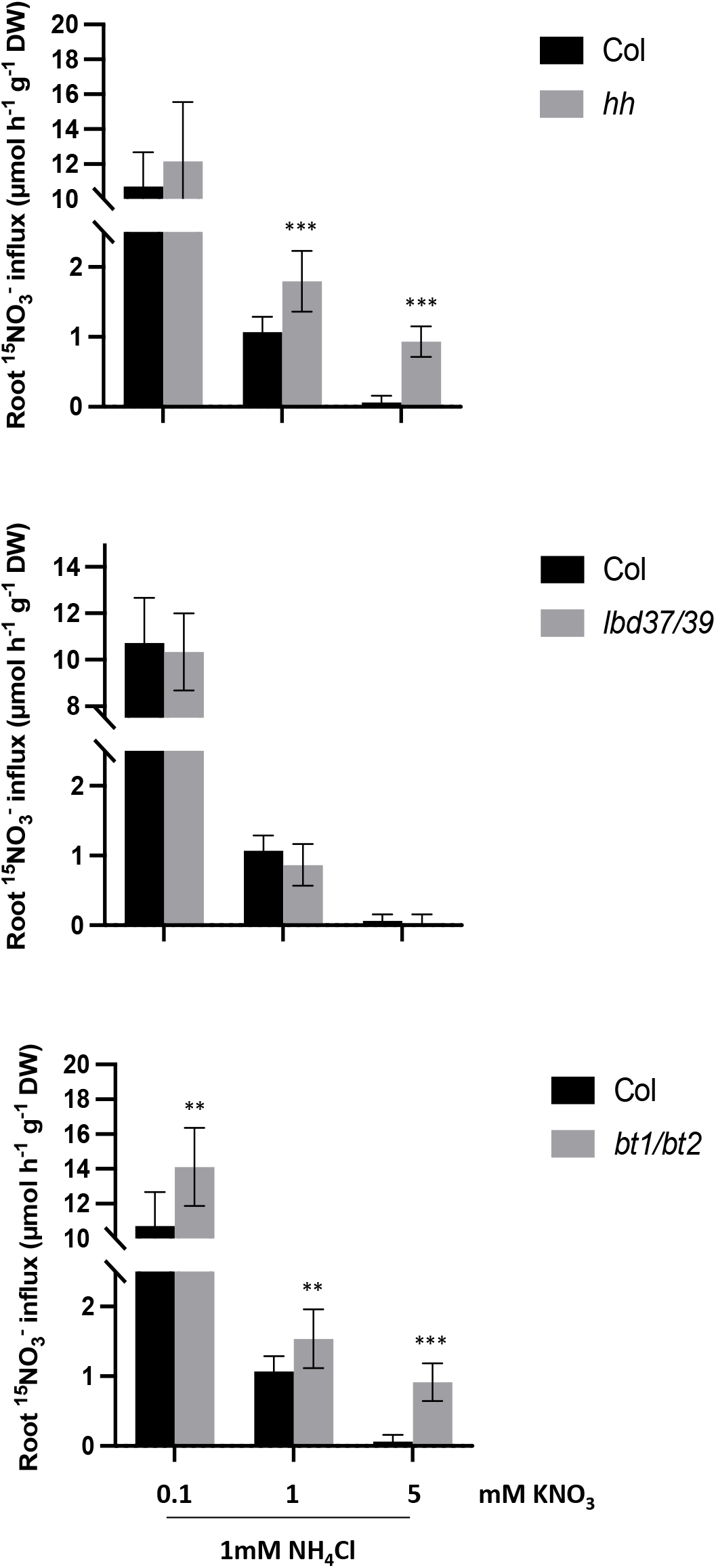
Impact of *hh (hho1/hrs1), lbd37/39* and *bt1/bt2* mutations on root NO_3_^-^ influx regulation by NO_3_^-^. Plants were grown on 1 mM NH_4_NO_3_ for 5 weeks before being transferred during 72h on 1 mM NH_4_Cl with 0.1, 1 or 5 mM KNO_3_. Root NO_3_^-^ influx was measured at the external concentration of 10 μM ^15^NO_3_^-^. Values are means of 12 replicates ± SD. Differences between WT (Col) and the mutants are significant at \**P* < 0.05, \*\**P* < 0.01, \*\*\**P* < 0.001 (Student’s *t* test).

**Figure 7.**
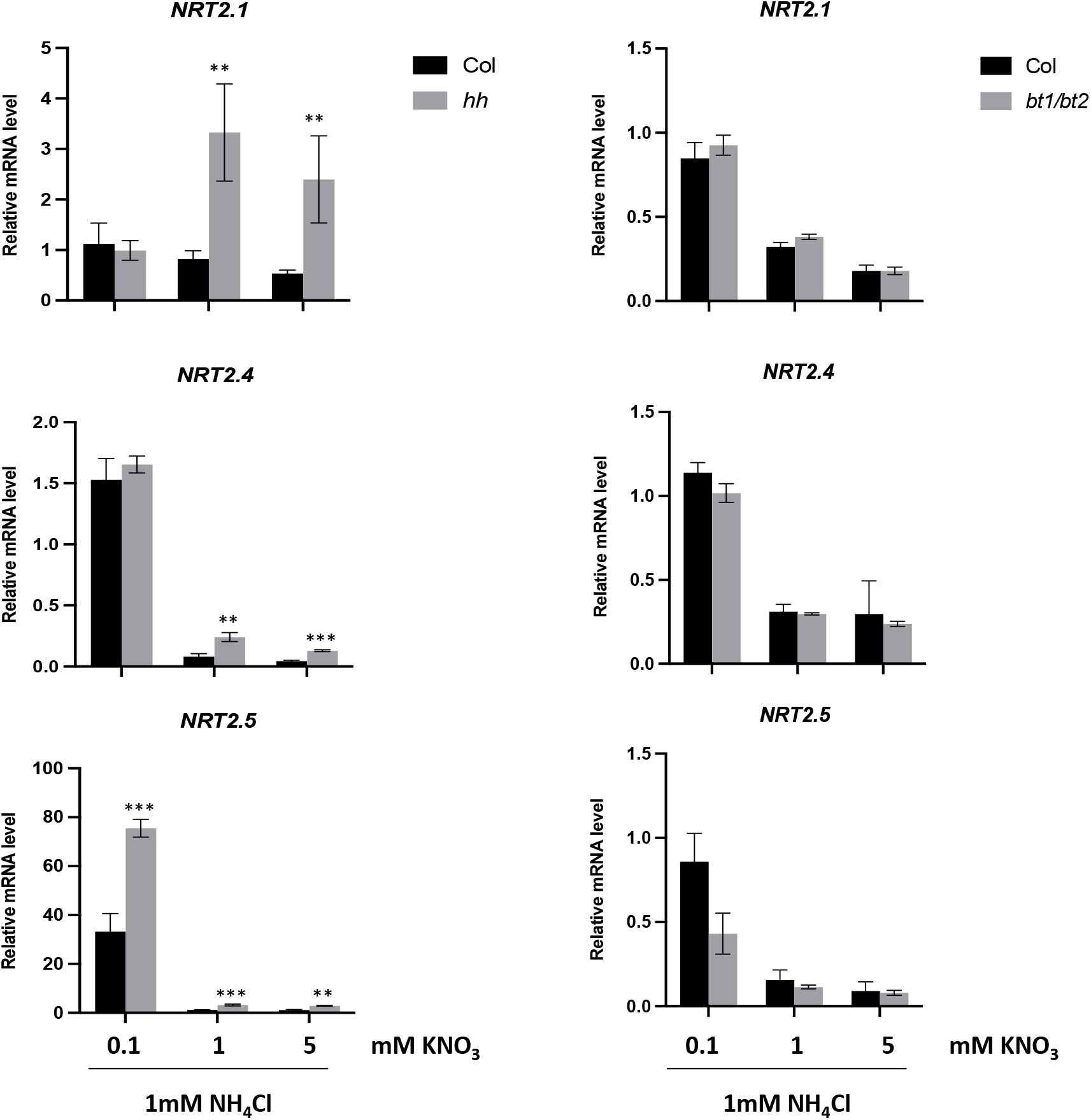
Impact of the double mutation *hho1/hrs1 (hh*) and *bt1/bt2* on *NRT2.1, 2.4, 2.5* repression by NO_3_^-^. Plants were grown on 1 mM NH_4_NO_3_ for 5 weeks before being transferred during 72h on 1 mM NH_4_Cl with 0.1, 1 or 5 mM KNO_3_. Roots have been collected to assess *NRT2.1, NRT2.4* and *NRT2.5* mRNA accumulation by RT-QPCR (relative accumulation to *Clathrin* housekeeping gene). Values are means of three biological replicates ± SD. Differences between WT (Col) and the mutant *chl1.5* are significant at \**P* < 0.05, \*\**P* < 0.01, \*\*\**P* < 0.001 (Student’s *t* test).

## Discussion

### *NRT2.4* and *NRT2.5* are repressed by NO_3_^-^ but not by N metabolites

Previous studies showed that *NRT2.4* and *NRT2.5* are, like *NRT2.1*, upregulated by N starvation (Lejay et al., 1999; Kiba et al., 2012; Lezhneva et al., 2014). Nevertheless, these N starvation experiments, consisting in transferring the plants from a growing solution rich in NO_3_^-^ to a solution with no N, did not allow to determine if NO_3_^-^ itself or N metabolites were involved in their regulation (Kiba et al., 2012; Lezhneva et al., 2014). For *NRT2.1*, it has been shown that both, N metabolites from NO_3_^-^ assimilation and high external concentration of NO_3_^-^ itself, are able to repress its expression (Lejay et al., 1999; Krouk et al., 2006). Our results show that this does not hold true for *NRT2.4* and *NRT2.5*, which appeared to be only repressed by high NO_3_^-^. Indeed, preventing normal NO_3_^-^ assimilation by NR knock-down in *g’4.3* plants failed to increase *NRT2.4* and *NRT2.5* expression compared to WT plants (Figure 1). Moreover, transfer of WT plants from NO_3_^-^ to NH_4_^+^ as sole N source dramatically increased the expression of *NRT2.4* and *NRT2.5*, while it repressed *NRT2.1* as previously observed (Figure 2) (Lejay et al., 1999). The repressive role of NO_3_^-^ for the regulation of *NRT2.4* and *NRT2.5* was confirmed by transferring the plants from 1 mM NH_4_NO_3_ to a solution containing 1 mM NH_4_^+^ but with increasing concentration of NO_3_^-^ (Figure 3A). This experimental protocol has been used previously by Krouk et al. (2006) to reveal the specific role of NO_3_^-^ in the repression of *NRT2.1*. Indeed, in those conditions, despite the continuous presence of NH_4_^+^, *NRT2.1* expression was consistently found to be determined by external NO_3_^-^ concentration, with a strong down-regulation as soon as NO_3_^-^ concentration exceeded 0.2 to 0.5 mM range (Krouk et al., 2006). Furthermore, using NRT1.1/NPF6.3 mutants, it was shown that *NRT2.1* repression by NO_3_^-^ was triggered by NRT1.1/NPF6.3. Our results show that this is also the case for both *NRT2.4* and *NRT2.5* that are, like *NRT2.1*, repressed by the increasing concentration of NO_3_^-^ in the presence of NH_4_^+^ in WT plants, but not in the *chl1-5* mutant (Figure 3A). In WT plants, the repressive effect of NO_3_^-^ was even stronger for *NRT2.5*, which was already downregulated by 0.1 mM of NO_3_^-^ compared to *NRT2.1* and *NRT2.4*. This is consistent with previous results showing that *NRT2.5* is not induced by NO_3_^-^ compared to *NRT2.1* and *NRT2.4* (Kotur and Glass, 2015). At the concentration of 0.1 mM NO_3_^-^ it is thus likely that, for *NRT2.1* and *NRT2.4*, the inductive effect of NO_3_^-^ overcome its repressive effect.

Bouguyon et al. (2015) suggested that in plants grown on 10 mM NH_4_NO_3_, repression of *NRT2.1* by high NO_3_^-^ is mediated by NRT1.1/NPF6.3 phosphorylated form on T101 residue. The results we obtained with our experimental setup confirmed this conclusion but also showed that this is not totally the case for both *NRT2.4* and *NRT2.5* (Figure 3B). Indeed, for *NRT2.1*, inhibition of T101 phosphorylation resulted in a complete lack of NO_3_^-^ repression on 1 mM and 5 mM NO_3_^-^, while for *NRT2.4* and *NRT2.5* the effect was only partial as both genes were still significantly downregulated by increasing NO_3_^-^ concentration in the T101A mutant plants. This suggests that the non-phosphorylated form of NRT1.1/NPF6.3 is somehow also able to mediate repression of *NRT2.1* by high NO_3_^-^, although less efficiently than the phosphorylated form. Compared with the results obtained with *chl1-5* mutant, it suggests that the regulatory mechanism triggered by NRT1.1/NPF6.3 is more complex and does not only depend on T101 phosphorylation for the repression by NO_3_^-^ of *NRT2.4* and *NRT2.5*.

In addition, our data indicate that NO_3_^-^ repression of *NRT2.4* and *NRT2.5* plays a key role in the regulation of very high affinity root NO_3_^-^ uptake. Indeed, ^15^NO_3_^-^ influx measurements at either 50 or 10 μM revealed that, in experiments where *NRT2.1* was not regulated like *NRT2.4* and *NRT2.5*, influx at 50 μM of ^15^NO_3_^-^ was correlated with *NRT2.1* expression, while influx at 10 uM was correlated with *NRT2.4* and *NRT2.5* expression (Figure 2A and 2B). This is in agreement with the role of very high affinity root NO_3_^-^ transporters attributed to both *NRT2.4* and *NRT2.5* (Kiba et al., 2012; Lezhneva et al., 2014).

### Repression of root NO_3_^-^ uptake by NO_3_^-^ involves key regulators of NRT2s both at the transcriptional and post-translational level

Analysis of the transcriptomic experiments performed by Bouguyon et al. (2015) revealed that several known repressors of *NRT2.1, NRT2.4* and *NRT2.5* are induced by 10 mM NH_4_NO_3_ and that this regulation is triggered by NRT1.1/NP6.3 T101 phosphorylated form (Supplemental Figure 1). Interestingly, a protein phosphatase from PP2C family was also found co-regulated with *NRT2.1* and *NRT2.4*. This phosphatase has recently been involved in the activation of NRT2.1 by directly dephosphorylating S501, a residue that functions as a negative phospho-switch in Arabidopsis (Jacquot et al., 2020; Ohkubo et al., 2021). Using our experimental set up with increasing concentrations of NO_3_^-^ in the presence of 1 mM NH_4_^+^ in both WT plants and the *chl1-5* mutant, we found that 7 out of the 10 regulators tested were regulated by high NO_3_^-^ in a NRT1.1/NPF6.3 dependent manner (Figure 4). Among them, we confirmed the results obtained by Bouguyon et al. (2015) for the regulation of *HHO1, HRS1, LBD39, BT2* and *PP2C*. However, in our hands *HHO3* was not found induced by high NO_3_^-^ in WT plants nor dependent on NRT1.1/NPF6.3 in the *chl1-5* mutant. Conversely, our results show that *LBD37* and *BT1*, which were not selected by Bouguyon et al. (2015), are both induced by NO_3_^-^ and dependent on NRT1.1/NPF6.3 signaling pathway (Figure 4). These discrepancies could be explained by the very different conditions between the experiments of Bouguyon et al. (2015) and ours, and suggest that the various members of the *HHO/HRS/NIGT* and *LBD* families are differentially regulated. A difference in the regulation of *HHO1*, *HRS1* and *HHO2*, *HHO3* has already been described by Kiba et al. (2018). In that case, it has been shown that *HHO2* and *HHO3*, unlike *HHO1* and *HRS1*, are induced by reduced forms of N such as Gln and urea. It supports the hypothesis that *HHO1* and *HRS1* are not involved in the same signaling pathways as *HHO2* and *HHO3*. Concerning LBDs and BTs, the work of Rubin et al. (2009) does not allow to identify different roles between LBD37, 38 et 39, while for BT1 and BT2 the work of Araus et al. (2016) indicate a functional redundancy suggesting that they are part of the same signaling pathway.

Surprisingly, despite the fact that *LBD37* and *LBD39* induction by NO_3_^-^ depends on NRT1.1/NPF6.3, it does not seem to specifically involve NRT1.1 phosphorylated form compared to the other molecular elements we identified (Figure 5). It supports the hypothesis, as discussed above, that the regulatory mechanisms triggered by NRT1.1/NPF6.3 are more complex and do not only depend on T101 phosphorylation for the repression by NO_3_^-^ of at least *NRT2.4* and *NRT2.5*. Although the double mutation of *LBD37/LBD39* had no effect of the repression of root NO_3_^-^ influx by high NO_3_^-^, those of *HHO1/HRS1* and *BT1/BT2* somehow attenuated it (Figure 6). Interestingly, if the impact of the double mutation of *HHO1* and *HRS1* on root NO_3_^-^ influx in *hh* mutant can be explained by the impact of these transcription factors on the expression of *NRT2s* transporters and especially of *NRT2.1*, this is not the case for BT1 and BT2 (Figure 7). It is surprising compared to previous results showing that on low NO_3_^-^, an increase of NO_3_^-^ uptake in *bt1/bt2* mutant was correlated with an increase in the expression of both *NRT2.1* and *NRT2.4* (Araus et al., 2016). However, once again, the experimental conditions were very different in Araus et al. (2016), with plants grown *in vitro* in steady state conditions with two different concentrations of NO_3_^-^. Furthermore, the molecular function of BT1 and BT2 proteins remains to be elucidated. Indeed, they are found in multisubunit E3 ubiquitin ligase complexes as well as in interaction with the BET10 transcriptional activator (Du and Poovaiah, 2004; Figueroa et al., 2005). It is thus possible that BT proteins are involved in both transcriptional regulation and/or degradation of proteins.

Finally, a particularly interesting result concerns the strong impact of NRT1.1 phosphorylated form on the repression by NO_3_^-^ of the protein phosphatase gene *PP2C* (Figure 5). This may help answering the unresolved question of the respective importance of transcriptional and posttranscriptional regulation of NRT2.1. On the one hand, changes in NO_3_^-^ HATS activity were always found highly correlated with changes in *NRT2.1* transcript accumulation and *NRT2.1* promoter activity in roots, suggesting a major role for transcriptional regulation (Lejay et al., 1999; Wirth et al., 2007; Girin et al., 2010; Laugier et al., 2012). This was further supported by the observations that the mutation or overexpression of key regulators governing *NRT2.1* transcription, such has NRT1.1, NLP7, HHO/HRS/NIGTs or CEPD/CEPDLs also resulted in a deregulation of the NO_3_^-^ HATS activity (Muñoz et al 2004, Yu et al. 2016, Kiba et al 2018, Maeda et al. 2018, Ota et al. 2020). On the other hand, suppression of the transcriptional regulation of *NRT2.1* using *35S* promoter failed to prevent feedback downregulation of the NO_3_^-^ HATS activity by N satiety or darkness, indicating a predominant role for posttranscriptional control (Laugier et al. 2012). Furthermore, several mechanisms have been proposed for posttranslational regulation of NRT2.1 (Wirth et al. 2007), among which phosphorylation of the S501 residue was shown to play a crucial role for governing HATS activity (Jacquot et al. 2020, Ohkubo et al. 2021). Our results showing that both *NRT2.1* and *PP2C* are common targets of the NRT1.1/NPF6.3-mediated repression of gene expression by high NO_3_^-^ allow to reconcile the above apparently contradictory observations. Indeed, this suggests that transcriptional regulation of *NRT2.1 per se* does not play a predominant role, but that the signalling pathways triggering this regulation are of crucial importance for controlling NO_3_^-^ HATS, because they also govern the expression of posttranslational regulators of NRT2.1 (Figure 3, Figure 4 and Figure 5, Ohkubo et al. 2021). Furthermore, this co-regulation of *NRT2.1* gene expression and NRT2.1 protein activity, through the regulation of the protein phosphatase gene *PP2C* does not only concern the repression by NO_3_^-^ since Ohkubo et al. (2021) showed that *PP2C* was regulated like *NRT2.1* in response to NH_4_^+^, NO_3_^-^ concentration in the media and N starvation. Altogether, these results suggest that the regulation of NO_3_^-^ HATS activity is the result of a redundant regulation of NRT2.1 at the transcriptional and post-translational level. It is interesting to note that redundant regulation at the transcriptional and post-translational level seems to be a general feature of the enzymes involved in N metabolism since it has already been described in plants for Nitrate Reductase (NR), Nitrite Reductase (NiR) and Glutamine synthetase (GS) (Crete et al., 1997; Campbell, 1999; Oliveira and Coruzzi, 1999).

## Conclusion

Altogether, as shown in Figure 8, our results allow us to propose a model for the signaling pathway downstream of NRT1.1 phosphorylated form and involved in high NO_3_^-^ repression of root NO_3_^-^ uptake. It involves the transcription factors HHO1 and HRS1, the proteins BT1 and BT2 and the phosphatase PP2C At4g32950. It revealed a complex picture, in which different level of regulation at the transcriptional and post-translational level are involved. HHO1 and HRS1 seem to be directly involved in the transcriptional repression of *NRT2.1, NRT2.4* and *NRT2.5* and this is supported by previous results showing that these two transcription factors can bind at least *NRT2.4* and *NRT2.5* promoter (Kiba et al., 2018; Safi et al., 2021). However, based on the results obtained with the mutants for NRT1.1 phosphorylated form, it seems that, for *NRT2.4* and *NRT2.5*, other elements are still missing to fully explained their repression by high NO_3_^-^. In the meantime, as shown for NRT2.1, the repression by high NO_3_^-^ of the PP2C protein phosphatase At4g32950 leads to an increase in the inactive form of NRT2.1 phosphorylated on S501. It revealed that the regulation of root NO_3_^-^ uptake in response to high NO_3_^-^ is likely the result of both a repression of NO_3_^-^ transporters at the transcriptional level and an inactivation at the protein level. Finally, despite the fact that BT1 and BT2 are involved in the repression of root NO_3_^-^ uptake by high NO_3_^-^, the molecular function of these proteins remains to be addressed.

**Figure 8.**
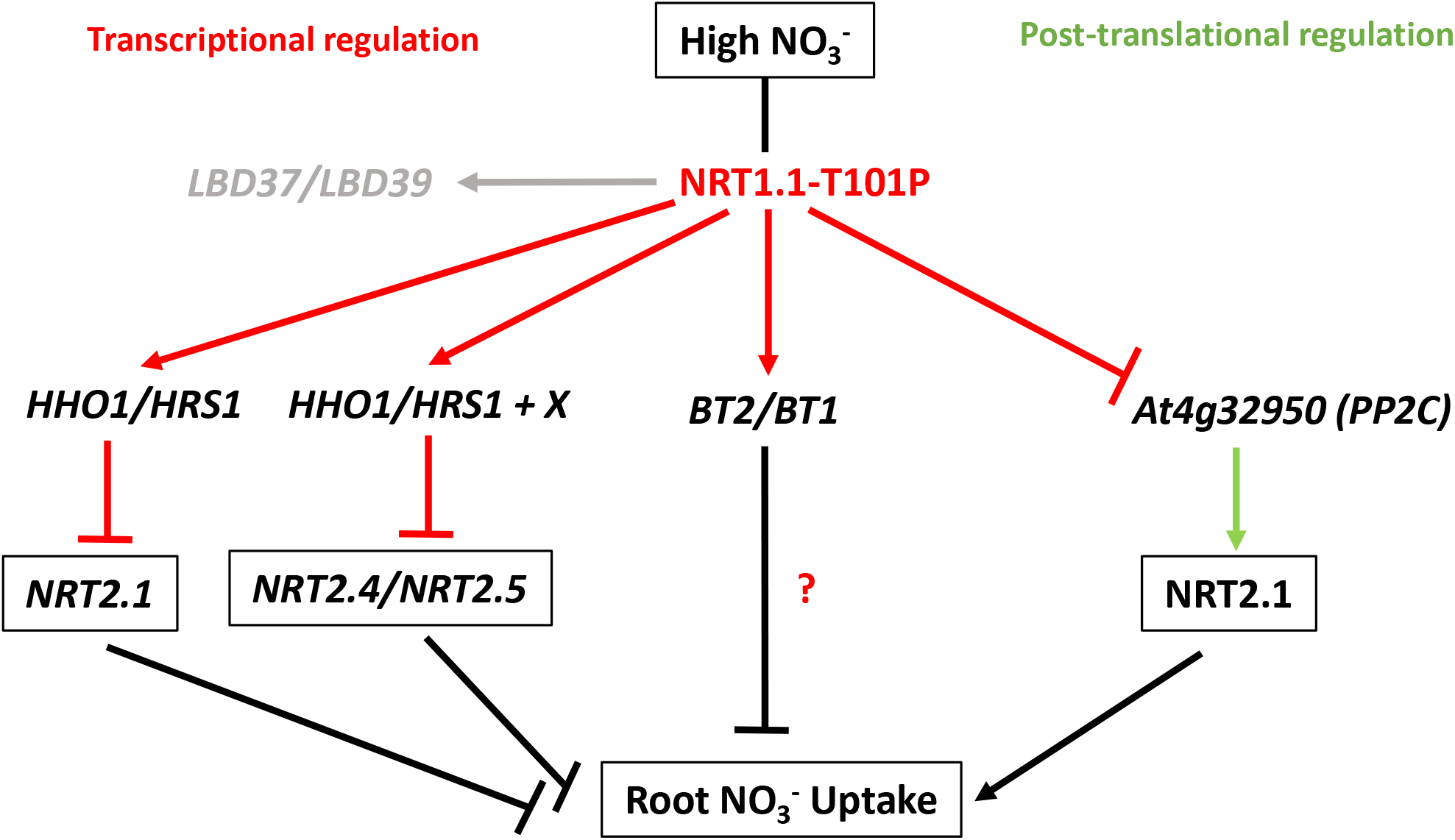
Model for the regulation of root NO_3_^-^ uptake by high NO_3_^-^ concentration

## Materials and Methods

### Plant Material

*Arabidopsis thaliana* genotypes used in this study were the wild-type Col-0 ecotype and the mutants *chl1-5* (Tsay et al., 1993), *g’4.3* (Wilkinson and Crawford, 1993), *hho1/hrs1 (hh*) (Medici et al., 2015), *bt1/bt2* (Sato et al., 2017) and *lbd37/lbd39 (lbd37-1*, SALK_097991; *lbd39-1*, SALK_049910).

In all experiments plants were grown hydroponically under non sterile conditions as described by Lejay et al. (1999). Briefly, the seeds were germinated directly on top of modified Eppendorf tubes filled with pre-wetted sand. The tubes were then positioned on floating rafts and transferred to tap water in a growth chamber under the following environmental conditions: light/dark cycle of 8 h/16 h, light intensity of 250 μmol·m^-2^·s^-1^, temperature of 22/20°C, and RH of 70%. After 1 week, the tap water was replaced with a complete nutrient solution containing 1 mM KH_2_PO_4_, 1 mM MgSO_4_, 0.25 mM K_2_SO_4_, 0.25 mM CaCl_2_, 0.1 mM FeNa-EDTA, 50 *μ*M KCl, 30 *μ*M H_3_BO_3_, 5 *μ*M MnSO_4_, 1 *μ*M ZnSO_4_, 1 *μ*M CuSO_4_, and 0.1 *μ*M (NH_4_)_6_Mo_7_O_24_. For growth of the plants, 1 mM NH_4_NO_3_ or 1 mM NO_3_^-^ was added to the medium as the N source as indicated in the text and figures. The plants were allowed to grow for 3 additional weeks before the experiments. Nutrient solutions were renewed weekly and on the day before the experiments. Depending on the experiments, 1 mM NH_4_NO_3_ or 1 mM NO_3_^-^ was replaced as a N source by either KNO_3_ or NH_4_Cl, or mixtures of these salts, as indicated in the text and figures.

### RNA Extraction and Gene Expression Analysis

Root samples were frozen in liquid N_2_ in 2-mL tubes containing one steel bead (2.5 mm diameter). Tissues were disrupted for 1 min at 30 s^-1^ in a Retsch mixer mill MM301 homogenizer (Retsch, Haan, Germany). Total RNA was extracted from tissues using TRIzol reagent (Invitrogen, Carlsbad, CA, USA). Subsequently 2 μg of RNA were used to perform reverse transcription in the presence of Moloney murine leukemia virus reverse transcriptase (Promega, Madison, WI, USA) after annealing with an anchored oligo(dT)_18_ primer as described by (Wirth et al., 2007). The quality of the cDNA was verified by PCR using specific primers spanning an intron in the gene *APTR* (At1g27450) forward 5’-CGCTTCTTCTCGACACTGAG-3’; reverse 5’-CAGGTAGCTTCTTGGGCTTC-3’.

Gene expression was determined by quantitative real-time PCR (qRT-PCR; LightCycler 480, Roche Diagnostics, Rotkreuz, Switzerland) using SYBR Premix Ex Taq™ (TaKaRa, Kusatsu, Japan) according to the manufacturer’s instructions. Conditions of amplifications were performed as described by Wirth *et al*. (2007), except the first 10 min at 95°C was changed to 30 s. All the results presented were standardized using the housekeeping gene Clathrin (At4g24550). Gene-specific primer sequences were: NRT2.1 forward, 5’-AACAAGGGCTAACGTGGATG-3’; NRT2.1 reverse, 5’-CTGCTTCTCCTGCTCATTCC-3’; NRT2.4 forward, 5’-GAACAAGGGCTGACATGGAT-3’; NRT2.4 reverse, 5’-GCTTCTCGGTCTCTGTCCAC-3’; NRT2.5 forward, 5’-TGTGGACCCTCTTCCAAAAA-3’; NRT2.5 reverse, 5’-TTTGGGGATGAGTCGTTGTGG-3’; HHO1/NIGT1.3 forward, 5’-GTAGGAAGATTTCGGAAGATAGAT-3’; HHO1/NIGT1.3 reverse, 5’-TTTGTACGAGTAGAACAAGACATAG-3’; HHO2/NIGT1.2 forward, 5’-AAACCAAAAAGCGGTGCGTT-3’; HHO2/NIGT1.2 reverse, 5’-ACTAGCTACTTTCACCGCCG-3’; HHO3/NIGT1.1 forward, 5’-ACTAATAATAGAGTTTACGCTCCTG-3’; HHO3/NIGT1.1 reverse, 5’-GGTGTGTGTGTAGTAGTAGAAGATG-3’; HRS1/NIGT1.4 forward, 5’-TTATAGACCGTCGATTATTGTGGA-3’; HRS1/NIGT1.4 reverse, 5’-TAATGATTACGGGTAGAAGAAGAC-3’; LBD37 forward, 5’-TGGATTGAAACCGCCGATGCTC-3’; LBD37 reverse, 5’-CGACTGAAACAAAGCAGGACGTTG-3’; LBD38 forward, 5’-TCAATGCCCTGCTTTGTTTCAGTC-3’; LBD38 reverse, 5’-AACCGCCGCTTGACAAACATTC-3’; LBD39 forward, 5’-CCTGAACTCCAACGTCCTGCTTTG-3’; LBD39 reverse, 5’-TTGGCATACGTGCCAGTTCCTG-3’; BT1 forward, 5’-CCGTTGAACAGACAGAAGGA-3’; BT1 reverse, 5’-CTGCATCGTCGATGAATTGG-3’; BT2 forward, 5’-TCCATTCGCAGTTTAAGACC-3’; BT2 reverse, 5’-AACTGGAGAATGTCGAGCTC-3’; PP2C forward, 5’-TGCTGTTCTCGCCGTTAAA-3’; PP2C reverse, 5’-TCCATCCTCACTTGTTCCAATC-3’; Clathrin forward, 5’-AGCATACACTGCGTGCAAAG-3’; Clathrin reverse, 5’-TCGCCTGTGTCACATATCTC-3’.

### NO_3_^-^ influx studies

Root NO_3_^-^ influx was assayed as described by (Delhon et al., 1995). Briefly, the plants were sequentially transferred to 0.1 mM CaSO_4_ for 1 min, to a complete nutrient solution, pH 5.8, containing 0.05 mM or 0.01 mM ^15^NO_3_^-^ (99 atom % excess^15^N) for 5 min, and finally to 0.1 mM CaSO_4_ for 1 min. Roots were then separated from shoots, and the organs dried at 70 °C for 48 h. After determination of their dry weight, the samples were analyzed for total nitrogen and atom % ^15^N using a continuous flow isotope ratio mass spectrometer coupled with a C/N elemental analyzer (model Euroflash Eurovector, Pavia Italy) as described in (Clarkson, 1986).

## Acknowledgments

We thank Dr. Gabriel Krouk and Dr. Anna Medicis for providing the seeds for the double mutant *HHO1/HRS1* (*hh*) and Dr. Shuichi Yanagisawa for providing the seeds for the double mutant *BT1/BT2 (bt1/bt2*).

---

**Supplemental Figure 1.** Candidate genes affected by *nrt1.1* mutation in *chl1.5* mutant in response to high N. Transcriptomic analysis of WT plants (Col) and *nrt1.1* mutants (*chl1.5, chl1.9, chl12, T101A and T101D*) grown *in vitro* during 8 days on 10 mM NH_4_NO_3_ (Bouguyon *et al*., 2015).

